# Spontaneous otocoherence of the active ear

**DOI:** 10.1101/2025.11.14.687084

**Authors:** Seth N.S. Peacock, Václav Vencovský, Rebecca E. Whiley, Natasha Mhatre, Christopher Bergevin

**Author notes:** **Competing Interest Statement:** None.

## Abstract

Spontaneous otoacoustic emission (SOAE) provides compelling evidence of active force generation inside the inner ear, although there is significant debate about the underlying generation mechanism(s). SOAE is commonly characterized by peaks in a spectral domain representation (as derived from a discrete Fourier transform), occurring at idiosyncratic frequencies unique to a given ear. Such is typically computed as an averaged magnitude spectrum that discards phase information. Here, we explore the hypothesis that SOAE phase, readily extracted from pre-existing recordings, contains complementary information. We propose several measures to use this information to quantify otocoherence (a form of autocoherence referring to a phenomenon of the ear), primarily by measuring the consistency in SOAE phase accumulation over a particular timescale. We present results based on recordings from different species with disparate inner ear morphologies (humans, barn owls, lizards). For regions of SOAE activity we extract time constants representing the timescale over which otocoherence is maintained. We demonstrate that these vary significantly across species and (for the barn owl and Tokay gecko where this data is available) appear to correlate with measures of auditory nerve fiber tuning. Additionally, we adapted the method to identify regions of weak SOAE activity among fluctuations in the noise floor. These methods can readily be employed to re-analyze SOAE waveforms previously collected from a variety of species, making them of broader comparative utility to reveal information about SOAE generation and thereby active cochlear mechanics.

**Significance Statement:** We examined spontaneous otoacoustic emission, a widely accepted hallmark of active auditory biomechanics, and describe a novel approach to extract measures of its self-coherence. This was explored comparatively across different species of disparate inner ear morphology to provide insight into how active amplification operates in the presence of noise. Our approach elucidates the notion that what has been traditionally referred to as “spontaneous activity” is not unstructured noise, but fluctuations that reveal internal system dynamics. The methodology described here is straightforward to employ and can be applied to a variety of data types, both within and beyond auditory neuroscience.

## Introduction

The vertebrate inner ear is an active sensor that uses metabolic energy to boost its ability to encode sound [Ashmore et al., 2010], allowing it to function as a subatomic displacement sensor [see Bergevin et al., 2025A]. As direct intracochlear physiology poses myriad challenges, one pertinent branch of empirical evidence characterizing the active ear is the observation of spontaneous otoacoustic emission (SOAE) [Hudspeth, 2008]. This faint sound, emitted in the absence of external stimuli and detectable using a sensitive microphone placed in the ear canal, show direct correlations to auditory perception such as threshold sensitivity [Long & Tubis, 1988]. SOAE is prevalent across the animal kingdom [e.g., Bergevin et al., 2015], including animal groups such as lizards, where morphological features considered crucial in coupling sensory cells, such as the tectorial membrane (TM), can vary dramatically [e.g., Manley, 2004; 2006].

While the biophysical mechanisms underlying SOAE generation remain controversial [e.g., Shera, 2022; Bergevin et al., 2025B], SOAE activity appears to be a by-product of the “cellular cooperativity” [Zweig, 2003] underlying the active processes powering the ear. Thus, there are strong motivations in auditory neuroscience to further elucidate precisely what SOAE reveals about peripheral auditory transduction.

Traditionally, SOAE is described as an idiosyncratic set of peaks in spectral magnitude (e.g., Fig.1A). These peaks can be relatively stable across timescales of the human lifespan [Abdala et al., 2017]. Analysis is commonly done in the spectral domain via the Fourier transform and predominantly employs “spectral averaging” over segments of the larger waveform to identify peaks [Penner et al., 1993]. Accordingly, only magnitudes (or their square) are considered and phase information is discarded or ignored, since including the raw phases will lead to destructive interference as the segments are averaged.

However, several lines of evidence suggest SOAE phase may not be useless. First, a pair of seminal studies [Bialek & Wit, 1984; Shera, 2003] developed SOAE analysis methods that included phase information and argued that such reveals important biophysical considerations (e.g., self-sustained oscillations in contrast to passively filtered noise). Second, a recent study simultaneously measured SOAE from both the left and right ears of a gecko [Roongthumskul et al., 2019]. Lizards commonly have an interaural canal that directly acoustically couples the inner surfaces of both tympanic membranes (i.e., eardrums); remarkably, phase information indicates that the two ears appear to synchronize via acoustic crosstalk in the middle ear space [Whiley et al., 2025]. Third, a study examining short time-scale fluctuations in filtered SOAE peaks, a process that intrinsically involves phase, showed that correlative behavior can readily exist both within a given peak and between peaks [Bergevin et al., 2025B]. Lastly, there are close relationships between SOAE and evoked OAEs [e.g., Zwicker & Schloth, 1984], indicative of the possibility that external sounds cause some form of entrainment of SOAE generators, thereby yielding useful phase-related information [e.g., SFOAE phase-gradient delays, used to estimate cochlear frequency selectivity; see Shera et al., 2002]. Despite these advancements, SOAE generation still remains poorly understood.

Taken together, we thereby consider the hypothesis that previously untapped information can be ascertained from SOAE phase extracted directly from time waveforms, obtained without any external stimulus. The key challenge is that the phase needs to be properly referenced. We describe several different approaches to essentially reference the signal to itself (e.g., via a time-delay, or a frequency shift) so to obtain a measure of self-coherence, or “autocoherence.” Our definition of autocoherence is biophysically motivated: the notion is that the underlying generators are acting in concert to produce the observed activity, consistent with related ideas such as “coherent reflection” [Zweig & Shera, 1995; Bergevin & Shera, 2010; Bergevin et al., 2015] and self-synchronization [Vilfan & Duke, 2008].

Using this method, we extract time constants representing the timescale over which self-coherence is maintained for a particular region of SOAE activity. These vary significantly across species and (for the barn owl and Tokay gecko where this data is available) appear to correlate with measures of tuning from auditory nerve fibers. We anticipate that this work and its comparative nature will inform/constrain models of the active ear, as well as provide a framework for previously recorded SOAE data to be meaningfully re-explored. Moreover, the approaches developed here may be extended to characterize other forms of spontaneous acoustic phenomena [Losi et al., 2025].

## Results

We provide a brief overview of the basic methodological approach here, with further details presented in the Methods. The key quantity we describe is a phase-centric self-coherence measure (*C*). There are two basic approaches to reference the phase. For the first, consider a series of short segments (*τ* seconds long) extracted from a much longer waveform. We compute the discrete Fourier transform (DFT) of each, providing us with phase estimates as a function of frequency. The phase in each frequency bin is then referenced to the phase of a nearby frequency bin (i.e., one is subtracted from the other), providing us with inter-frequency phase differences Δ*ϕ*_*ω*_ as a function of segment and frequency. The motivation is biophysical: if generators for a specific peak are distributed across some narrow frequency band (e.g., a”cluster” as in the model of Vilfan & Duke, 2008), then there may be meaningful timing information maintained across adjacent bins of the DFT. After wrapping these phase differences into the range [−*π, π*], we then take their average absolute value for a given frequency for a quantity we denote < |Δ*ϕ*_*ω*_| >. If the phases are random (i.e., uniformly distributed in [−*π, π*]), this quantity will be near *π*/2 (the baseline value for the measure). Values lower than this baseline at a particular frequency indicate that the phase differences tended to be closer to 0 (the frequency bin tended to be in phase with its neighbor), whereas values higher than this baseline indicate phase differences closer to *π* (an antiphase relationship).

For the second, the signal is self-referenced to a time-delayed version of itself. The motivation is that timing information intrinsic to the system persists over time; specifically, self-coherent oscillations will accumulate consistent amounts of phase over a given reference time. Consider two short consecutive time segments, each *τ* seconds long, where the phase is measured (across frequency bins) for each segment with the DFT. From these two phases, we compute the phase difference Δ*ϕ_τ_* from one segment to the next, representing the phase accumulation for that frequency bin over *τ* seconds. Rather than strictly using consecutive segments (i.e., a *τ*-length shift between segment start times), we can also measure the phase accumulation across two *τ* length segments whose start times are separated by *ξ* seconds (a symbol visually reminiscent of a cochlear spiral), which we denote Δ*ϕ*_*ξ*_. Note that if *ξ* < *τ*, there will be overlap between the two time segments, and that Δ*ϕ_τ_* is then a special case of Δ*ϕ*_*ξ*_. We can then average across phase differences obtained by repeating this process for pairs of segments throughout the waveform to compute < |Δ*ϕ*_*ξ*_ | > similarly to < |Δ*ϕ*_*ω*_| >.

Now, rather than taking the averaged absolute value of the phase, we can instead compute a vector strength measure [e.g., Roongthumskul et al., 2019]. Such directly quantifies the consistency of the phase differences in a way which respects their 2*π* periodicity and is agnostic to the particular value they may (or may not) cluster around. Quantifying the consistency of the phase accumulations over time in this way leads to 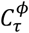 and 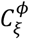 (again, the former being a special case of the latter where *ξ* = *τ*). For the analogous measure obtained from the frequency-referenced phases we use the symbol 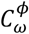. We refer to these as “autocoherence” measures in general, with the word “otocoherence” denoting the phenomenon in the ear quantified by this measure.

To visually demonstrate the method, Figure 1 shows the analysis of a waveform acquired from a green anole lizard (*Anolis carolinensis*). This choice of animal is motivated by the observation that the anole inner ear is relatively morphologically simple (e.g., short basilar membrane ∼0.35 mm long, ∼150 hair cells, no overlying TM for the majority of hair cells) yet demonstrates robust SOAE activity over the range of 1-7 kHz (depending upon body temperature) [Manley, 2006; Bergevin et al., 2025B]. Figure 1A shows a typical magnitude spectrum, where numerous distinct peaks of varying height and width are readily observed, as is a broader underlying “baseline” above the noise floor [Manley et al., 1996].

**Figure 1.**
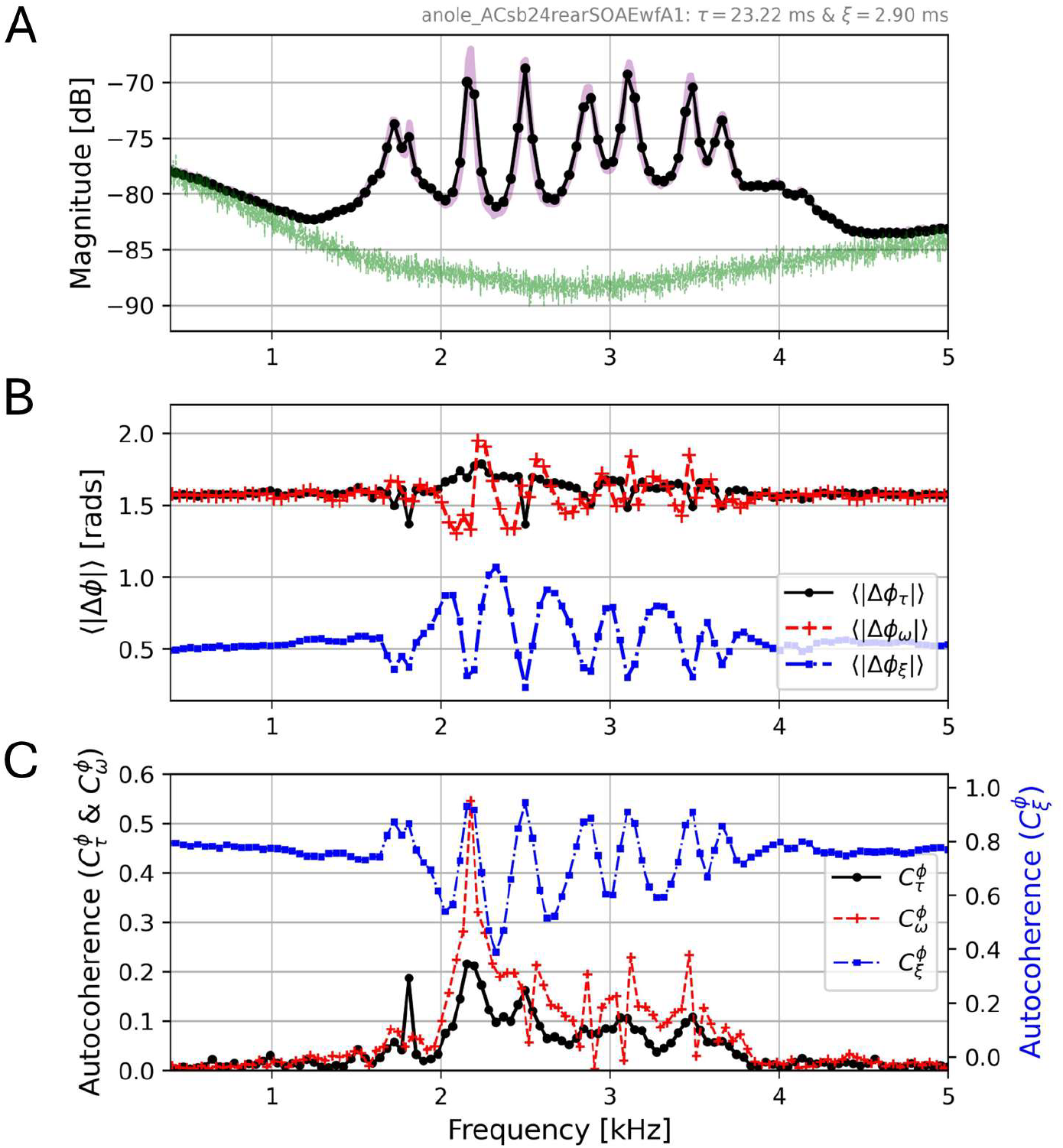
SOAE data from a representative green anole lizard (*Anolis carolinensis*). **A**: Traditional averaged magnitude spectrum obtained from a one-minute recording. The black curve is a time averaged magnitude spectrum using segments of length *τ* ≈ 23.2 ms (1024 samples) while the purple curve used segments of length *τ* = 80 ms (3528 samples) for improved frequency resolution. Also shown in green is the microphone noise floor. For an absolute reference, the noise floor at 3 kHz is approximately -20 dB SPL re. 2e-5 Pa. **B**: Only phase information was used to determine < |Δ*ϕ*| >. The three different curves represent different strategies for referencing the phase (time delayed by different durations in blue and black, frequency shifted in red). A reference time of *ξ* ≈ 2.9 ms (128 samples) was used for the blue curve, and all curves used segments of length *τ* ≈ 23.2 ms (1024 samples). Note the lower baseline in < |Δ*ϕ*_*ξ*_ | >, a manifestation of the “intrinsic gradient” as described in Methods. **C**: Autocoherence values obtained from the phase information alone. Note the dual ordinate axes and the elevated (blue) 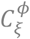 values, which occur for the same reason as the lower < |Δ*ϕ*_*ξ*_ | > and are the motivation for dynamic windowing (see Methods). All measures in this figure use a “boxcar” (aka no) window; also, a high pass filter (Kaiser design with a 150 Hz cutoff, 100 Hz transition region, and -100 dB allowed ripple) was applied to remove low frequency artifacts.

To first affirm our leading hypothesis, that SOAE phase alone contains information at least congruent to the magnitudes, Fig.1B shows various < |*Δϕ*| > measures. We see that in frequency regions where SOAE are absent, the phase averages to a uniform value. However, in regions of SOAE activity, deviations from this value are clearly present, and the largest excursions appear at frequencies close to the magnitude peaks. These curves therefore indicate that both approaches for phase referencing lead to some indication of SOAE activity distinct from the surrounding noise floor. However, these data are difficult to interpret given that both positive and negative deviations from ∼*π*/2 indicate a departure from random phases.

We further improve on this observation by instead calculating the vector strength-centric autocoherence. In the simplest measure, 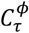 (black curve, Fig.1C), autocoherence values well above zero can be easily observed in frequency region of SOAE activity. If we use a shorter time delay (*ξ* < *τ*), 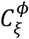 shows even more structure (blue curve, Fig.1C), albeit with an elevated baseline away from SOAE regions. This is due to the significant overlap between each pair segments used to calculate the phase differences Δ*ϕ*_*ξ*_: even for “noise bins,” the phase estimate from one segment will be highly correlated with the next, leading to spuriously high consistency of phase accumulations (see the “intrinsic gradient” described in Methods). Finally, the frequency-referenced 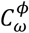 (red curve, Fig.1C) appears similar (e.g., similar peaks to 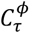 especially near magnitude peak frequencies), yet there are also notable differences. In summary, Fig.1 demonstrates that phase information alone can be used to characterize SOAE in a fashion which may be complementary to analyzing magnitudes alone.

We extend the same analysis to waveforms recorded from an individual human (Fig.2A), barn owl (Fig.2B), and a different green anole (Fig.2C) to characterize otocoherence across species and examine interspecific variation. All three species exhibit peaks corresponding to those apparent in the magnitudes, even upwards of 10 kHz. Note the time-delayed (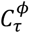 and 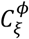) and the frequency-neighbor values 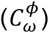 appear to yield complementary patterns; for example, 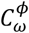 can be larger than 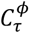 for some peaks and lower for others, though these differences were not systematically explored. An important difference between the 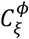 values in Fig.2 and those shown in Fig.1 is the lower (near zero) baseline values away from regions of SOAE activity. This was due to the larger *ξ* value used in Fig.2C compared to Fig.1, meaning less overlap between segments used to compute the phase accumulations Δ*ϕ_ξ_*.

**Figure 2.**
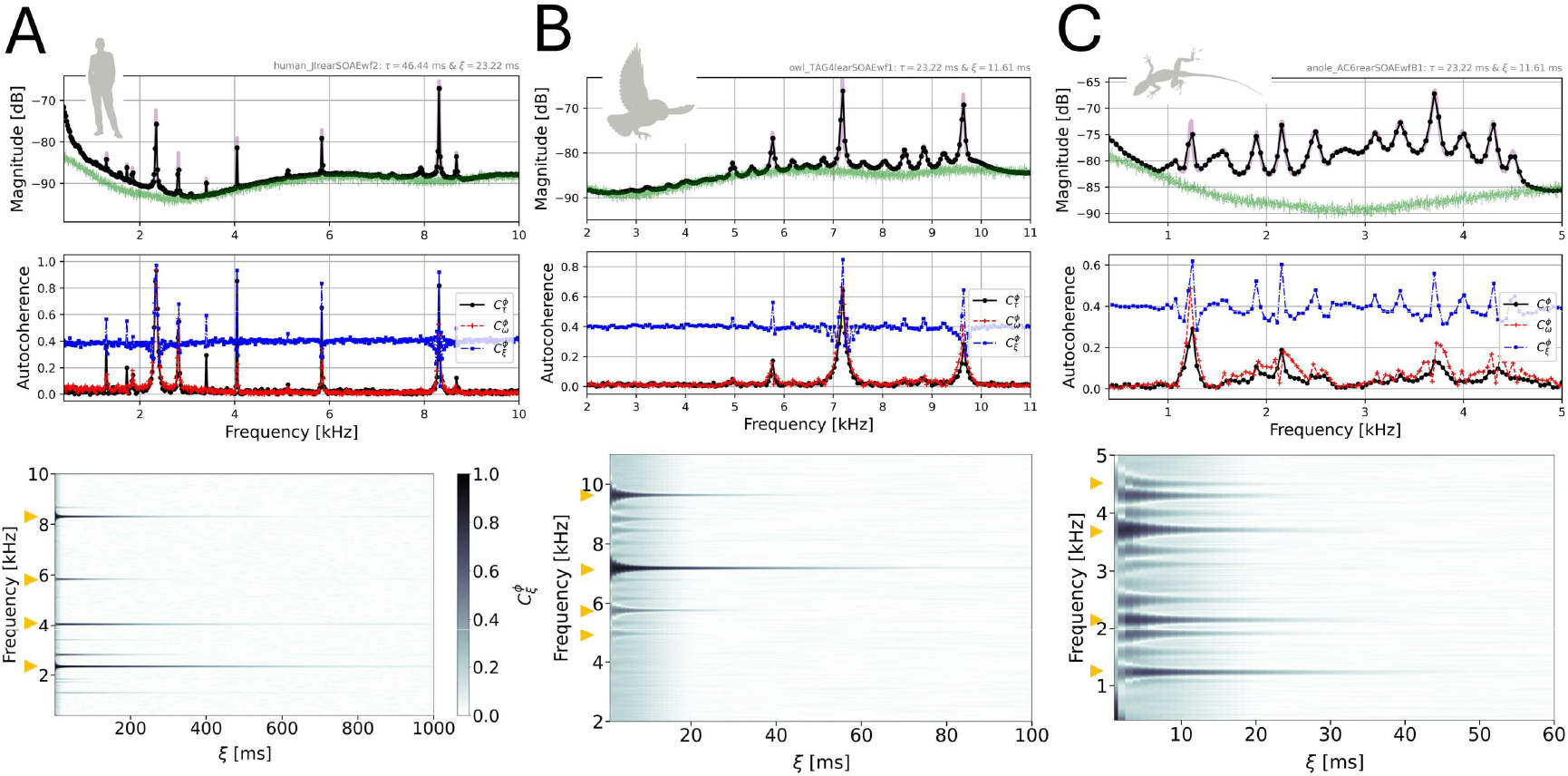
**Top row**: Spectral magnitudes and autocoherence values (similar to Fig.1) for a representative human (**A**), barn owl (**B**, *Tyto alba*), and green anole lizard (**C**, a different individual than the one shown in Fig.1). The same high pass filter described in Fig.1 was applied and the same *τ* = 80 ms (3528 samples) for the high resolution (purple) magnitudes; for the other measures, *τ* ≈ 46.4 ms (2048 samples) and *ξ* ≈ 23.2 ms (1024 samples) were used for the human, while *τ* ≈ 23.2 ms (1024 samples) and *ξ* ≈ 11.6 ms (512 samples) were used for the owl and anole. Note that larger *ξ* values were used in Fig.2C compared to Fig.1C; this had the effect of decreasing the baseline 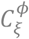 (blue) values due to a decrease in overlap between segments used to compute phase accumulations. **Bottom row**: The corresponding “colossograms” showing the decay of autocoherence with increasing lengths of *ξ* (using the parameters described in Methods for the *N_ξ_* analysis). Here, 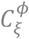 (see colorbar) is plotted as a function of both frequency and *ξ*. Note the different abscissa scales across the three columns. Small orange triangles in the colossograms indicate frequencies included in Fig.3 analyses.

The bottom row in Fig.2 introduces “colossograms,” from coherence-loss-o-gram, showing how the time-delayed autocoherence 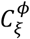 evolves with increasing lag values *ξ*. For these calculations, dynamic windowing was applied (see Methods). Briefly, each time segment was multiplied by a Gaussian window function whose width was a function of *ξ* (i.e., a small *ξ* would have a narrow width, to reduce the effects of the corresponding high degree of overlap). This allowed for distinguishing autocoherence decays over short *ξ* values which would otherwise be surrounded by an “intrinsic gradient” of uniform decay at all frequencies as the baseline decreased with increasing *ξ*. To see how this baseline depends on *ξ* in the absence of dynamic windowing, compare Fig.2C with Fig.1C. All autocoherence measures reported beyond Fig.2 used dynamic windowing. While this smooths the spectrum across frequency for low *ξ*, a key advantage of the dynamic approach is that it then gradually restores the desire frequency resolution for higher *ξ*. This restoration can be seen in how the high- 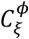 regions become narrower as *ξ* increases in the colossograms of Fig.2.

It was observed that human SOAE peaks maintained otocoherence almost an order of magnitude longer than those from owls or anoles. Peaks with longer persistence often also had higher magnitude, but not always (e.g., in Fig.2C the peak at ∼1.2 kHz persists appreciably longer than the one at ∼2.2kHz, but have nearly identical magnitudes).

A key aspect of the colossograms shown in Fig.2 is that each peak has a characteristic decay in 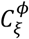. To explore this, consider Fig.3A, which shows the 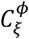 decay for two frequencies associated with SOAE peaks for a human and a green anole. For very short *ξ*, 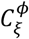 is small and increases from zero. This can in part be attributed to the dynamic windowing, since then small *ξ* means a narrow time window; the localization in time leads to wide frequency bins, each capturing a large range of oscillatory components whose collective phase will not accumulate consistently. After this initial rise, the autocoherence decays (see Fig.3A). To first order, the decays appear approximately exponential in nature. As such, we used nonlinear regression to estimate the associated exponential decay constant (*T*_*ξ*_). This time constant can then be multiplied by the peak’s center frequency for *N*_*ξ*_ = *T*_*ξ*_ ⋅ *f*_0_, which expresses the decay time in number of cycles. Figure 3B shows how *N*_*ξ*_/*π* varies across frequency for the three comparative species (humans, owls, and anoles), as well as one additional lizard species (Tokay gecko). For this plot, several arbitrarily chosen SOAE peaks were analyzed from at least four different individuals for each group. Broadly, the non-mammals share a similar trend, with *N*_*ξ*_ values increasing with frequency. However, the humans appear distinctly different, with *N*_*ξ*_ being nearly an order of magnitude larger.

**Figure 3.**
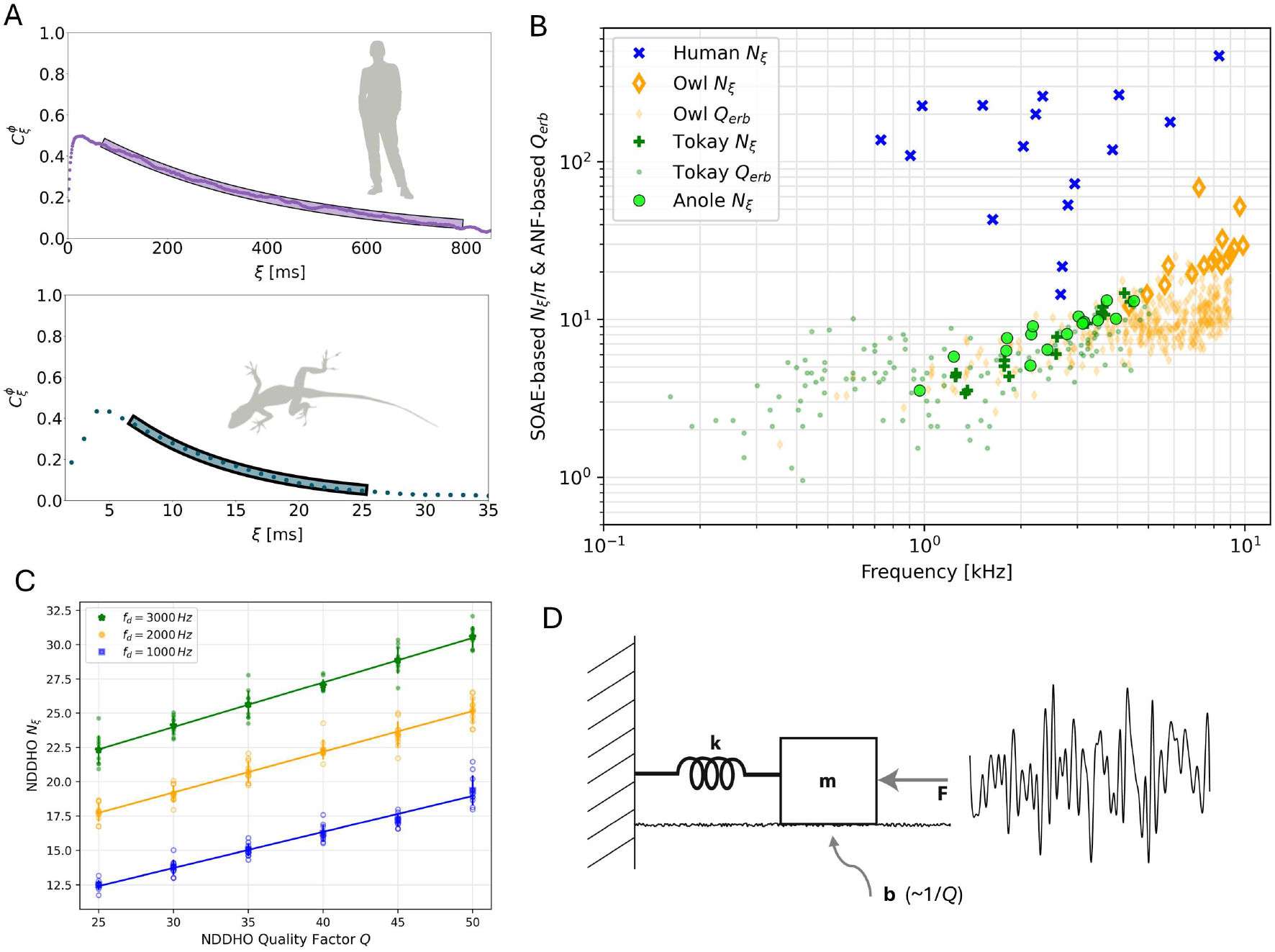
**A**: To demonstrate how *C*_*ξ*_ varied with increasing *ξ*, evaluated at the fixed center frequency of an SOAE peak (using the main analysis autocoherence parameters described in Methods), representative values (dots) are shown for both a human (0.904 kHz peak for subject TH13) and anole (4.50 kHz peak for lizard shown in Fig.2C). Note that for small *ξ* values, 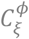 rises from zero and approaches a maximum value before starting to decrease. For that latter portion, an exponential was fit to the data (shaded region) via nonlinear regression to estimate the associated time constant *T*_*ξ*_. That value was then multiplied by the emission frequency to obtain *N*_*ξ*_, a decay time expressed in the number of oscillation cycles. For the two plots in panel A, the values of *N*_*ξ*_ were 343.0 (human) and 41.2 (anole). **B**: Two independent quantities are plotted together here. First, the *N*_*ξ*_/*π* is plotted for several peaks from four different individuals for each of the three comparative groups (human – blue crosses, barn owl – large/thick orange diamonds, green anole – large lime circles), as well as one additional group, the Tokay gecko (*Gekko gecko*, dark green pluses). Second, for the two species where single-unit auditory nerve fiber (ANF) threshold data have been reported (owl and gecko), the corresponding frequency selectivity expressed as *Q*_*ERB*_ is also plotted (small/thin orange diamonds and small dark green circles respectively). **C**: As shown by the schematic on the right side, the noise-driven damped harmonic oscillator (NDDHO) was numerically integrated and the associated *N*_*ξ*_ values extracted at the oscillator’s damped characteristic frequency (*f*_*d*_). Simulations were run for three different *f*_*d*_ values, with the quality factor (*Q*, inversely proportional to the damping coefficient) systematically varied. For each condition, ten different stochastic drives were simulated (transparent points) and the average and standard deviations plotted. A linear regression was also computed (straight lines).

A key finding of the present study emerges from Fig.3B by comparison of *N*_*ξ*_/*π* to measures of frequency selectivity obtained from single unit auditory nerve fibers (ANFs). We analyzed threshold tuning curves obtained from barn owl (Koppl, 1997) and Tokay gecko (Manley et al., 1999), which are the two groups where suitable ANF responses have been reported in addition to SOAE data. Specifically, we computed a *Q*_*ERB*_ values (*Q* is the quality factor and *ERB* stands for equivalent rectangular bandwidth) by integrating the area underneath an inverted tuning curve (see Bergevin et al., 2015). For reference, the commonly reported *Q*_10*dB*_ value is typically proportional to *Q*_*ERB*_ by a factor of ∼1.8. For both the owl and gecko, Fig.3B indicates that *N*_*ξ*_/*π* closely matches the upper bound of *Q*_*ERB*_ values. In short, the estimated time constant indicative of loss of otocoherence predicts the upper bound of frequency selectivity encoded into the central nervous system for both barn owl and Tokay gecko. This indicates that for ears exhibiting SOAE activity, the degree of cochlear tuning possible can readily be estimated from short non-invasive measurements without the issues and challenges associated with single-unit neural recordings (e.g., terminal experiments, presentation of multiple stimuli).

To further validate this observation relating tuning and autocoherence, we constructed a simple (linear) heuristic, the noise-driven damped harmonic oscillator (NDDHO, see Fig.3D). In analogy to the auditory filters of the inner ear, the DHO can act as a passive band-pass filter with a characteristic degree of tuning (as represented by the quality factor *Q*, which is inversely proportional to the amount of damping). We simulated the NDDHO numerically (see Methods), from which colossograms and decay constants could then be computed. We investigated how *N*_*ξ*_, here acting as a non-auditory measure of autocoherence, varied with respect to both the *Q* and the oscillator’s damped natural frequency *f*_*d*_. We found that for this linear filter, *N*_*ξ*_ and *Q* are proportional (Fig.3C). These results suggest that autocoherence inferred from the “spontaneous” responses can thereby be used to estimate the frequency selectivity of a system, and can be extended to the auditory system where some combination of structural and cellular tuning emerges in the presence of both internal (i.e., biological) and external sources of noise.

Lastly, we sought to explore whether the methods described here could be used to better detect the presence of SOAE peaks (e.g., small amplitude peaks close to the noise floor; see Talmadge et al., 1993). Detecting such peaks is important for validating different classes of models of SOAE generation, where interpeak spacing is a key consideration [e.g., Shera, 2003; Fruth et al., 2014]. Simply put, lowering the noise floor is desirable, but challenging for technical reasons such as Johnson–Nyquist noise related to the microphone and associated amplifier. Typically, “spectral averaging” helps reveal SOAE structure (e.g., Fig.1A), where the magnitudes of the Fourier transform (or their square, and hence power; see Methods) are averaged and the phase is ignored. This averaging smooths the noise floor but does not lower it.

Using time-referenced phase accumulations Δ*ϕ*_*ξ*_, we propose a new method: rather than simply averaging magnitudes, we average complex numbers whose phase is Δ*ϕ*_*ξ*_. These will add constructively if (and only if) the phase accumulations Δ*ϕ*_*ξ*_ are consistent. We develop a new metric denoted as 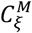, which utilizes both magnitude and phase information (see Eq.4), a spectrum with units of magnitude. Peaks in this spectrum then represent frequencies with both magnitude and otocoherence higher than their surroundings.

The top row of Fig.4 (panels A and B) shows examples of spectra for a Tokay gecko and a human with magnitude-only averaging (black) versus phase- and magnitude-based averaging 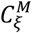 (blue). Overall, the structure of the peaks and valleys becomes more pronounced in 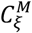, making it easier to determine if a peak is present and at what frequency. Our analysis therefore indicates this phase-centric averaging approach can be used to better characterize activity close to the noise floor. To further demonstrate this, we considered waveforms from 12 human subjects with robust SOAE activity and where interpeak spacing was computed non-dimensionally as *N*_*SOAE*_ (the geometric mean frequency of the pair divided by their difference; see Shera, 2003].

**Figure 4.**
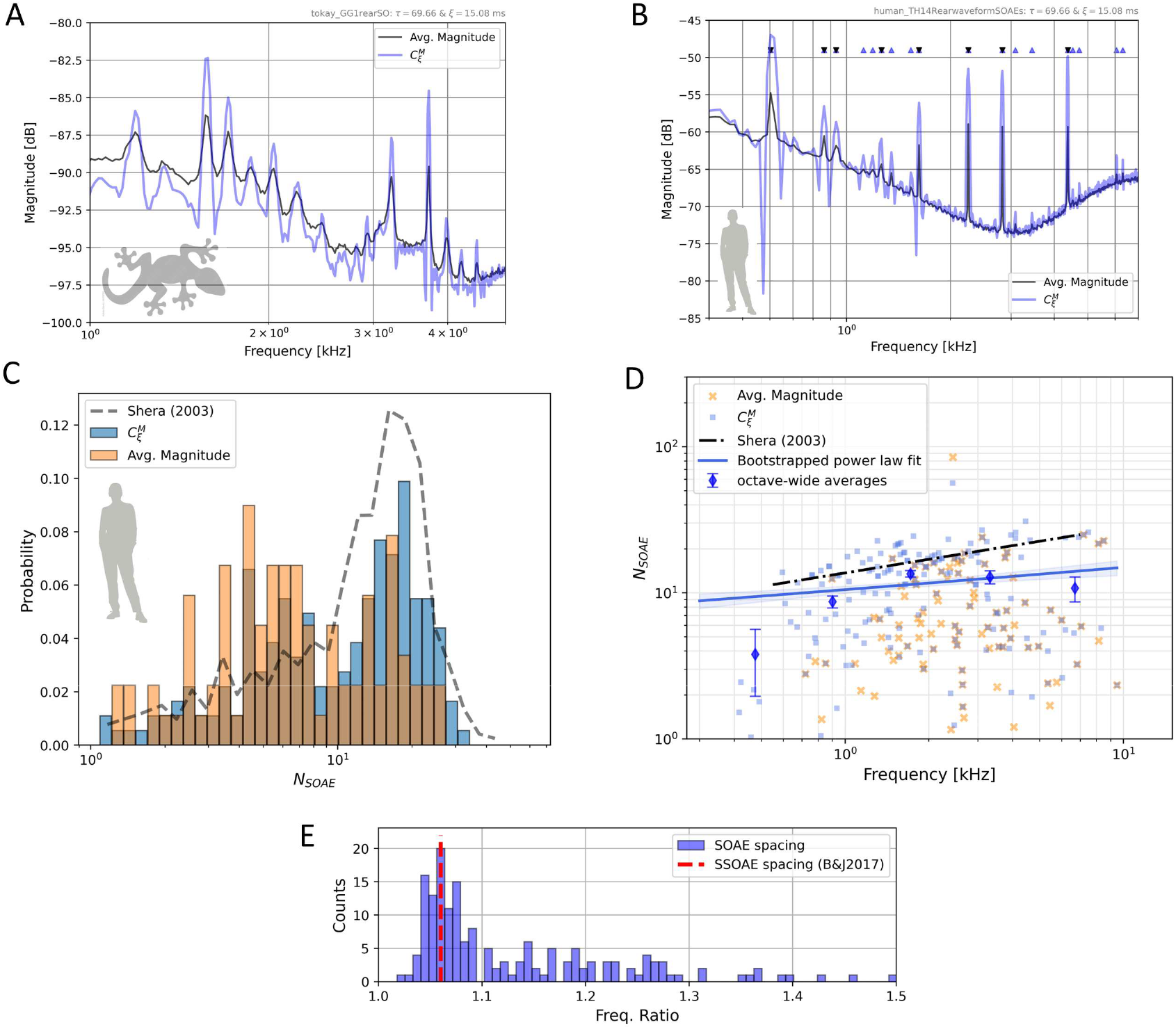
The top row (panels **A** and **B**) show the effects of two different averaging strategies, one for a representative Tokay gecko (Gekko gecko, **A**) and human (**B**). For each plot, averaged magnitudes (with a Hann window) are shown by the black curve and the autocoherence variant 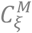 is shown in blue. For the human shown in panel B, SOAE peaks identified (see text) via the averaged magnitude approach are indicated by the downward black triangles, while peaks identified via 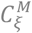 are noted by the upward blue triangles (there were 2.25 times more peak frequencies found). The parameters *τ* ≈ 69.66 ms (3072 samples) and *ξ* ≈ 15.08 ms (665 samples) were used. **C** – Distribution of *N*_*SOAE*_ values, defined as the ratio of the geometric mean frequency to the frequency difference between adjacent SOAE peaks. Data were compiled from 12 human subjects, all with relatively robust SOAE activity. Also included (dashed line) are the values reported by Shera (2003). **D** – Same data as shown in panel C, but the frequency-dependence is also shown. Similar to Shera (2003), a power law fit was computed (solid blue line) and bootstrapped to determine the associated uncertainty as a standard deviation (blue shaded area). Additionally, mean *N*_*SOAE*_ values for peaks identified via 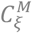 were computed in octave-wide bins (blue diamonds) and the associated standard error. **E** – Comparison of frequency ratio for adjacent SOAE peak pairs (identified via 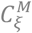, blue bars) compared to the 1.06 ratio (red dashed line) reported by Bell & Jedrzejczak (2017).

In both the averaged magnitude spectrum and 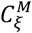, an SOAE peak was identified if it appeared by inspection to be an elevated region whose maximum was at least 2 dB above its surroundings. Many more peaks were identified in the 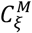 spectrum (blue upward triangles in Fig.4B as opposed to the black downward triangles marking peaks identified via averaged magnitudes). Across the dozen subjects, magnitude averaging revealed a total of 90 peaks, whereas 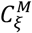 indicated 183 (or ∼15 peaks/ear), an increase of ∼103%. Since more peaks were detected, the overall interpeak spacing decreased, resulting in larger *N*_*SOAE*_ (Fig.4C displays the difference in *N*_*SOAE*_ distributions obtained via 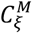 in blue and averaged magnitudes in orange) more in line with Shera (2003).

However, in contrast to that study, we found that the frequency dependence of *N*_*SOAE*_ was not well fit by a power-law dependence. Specifically, the bootstrapped range of power law fits do not align well with octave-wide bin averages (Fig.4D), nor with the reported fit from Shera (2003). We also observed that the most common interpeak ratio is close to 1.06 (Fig.4E), consistent with the value reported for synchronized spontaneous otoacoustic emissions (SSOAEs), where SOAE activity is stimulated by virtue of a low-level broadband stimulus [Bell & Jedrzejczak, 2017; see also Wable & Collet, 1994]. Those authors interestingly note that this ratio “suggests the human cochlea may be naturally configured to detect semitone intervals.”

## Discussion

### SOAE Autocoherence → Otocoherence

In this study, we developed a new method to extract information on SOAE phase stability that is relatively easy to implement. We showed that averaged phases contain spectral features that align with the region of SOAE activity, as inferred from magnitude information, and we developed several measures of otocoherence that exhibit quantitative similarities to sound-evoked neurophysiological measures. Aside from establishing that the entirety of SOAE waveforms contains meaningful biophysical information, how do these results progress what SOAE activity reveals about the active ear?

First, we examine the timescales over which SOAE self-coherence in phase is maintained (the colossograms of Fig.2). This allows us to probe the degree to which the oscillations are able to maintain consistency in the face of noise. Moreover, by using *ξ*-adjusted phases to improve the ability to detect peaks (Fig.4), we can reduce the effects of noise inherent to the measurement while isolating the signal emanating from the SOAE generation mechanism(s). While this SOAE activity is stochastic, it comes from an active system that is being driven/affected by noise [van Dijk et al., 1996; Shera, 2003]. Correspondingly, these otocoherence measures allow us to distill out those properties characteristic of that system.

Second, consider that there are close relationships between SOAE and emissions evoked by external sound stimuli (evoked otoacoustic emissions, or eOAEs) [e.g., Zwicker & Schloth, 1984; Shera, 2003; Bergevin et al., 2015]. These relationships potentially indicate that the external stimuli entrain SOAE generators, thereby yielding useful (stimulus-referenced) phase information that contributes to eOAE phase-derived quantities. For example, stimulus frequency emission (SFOAE) phase-gradient delays appear to correlate with cochlear tuning [Shera et al., 2001]. In turn, our data suggest that traces of eOAE-related phase behavior are intrinsically present even in the absence of any entraining stimuli (e.g., *self*-synchronization might be apparent in SOAE phase). A stimulus-independent measure circumvents the need for assumptions such as “low to moderate sound levels” [Shera et al., 2010] to address the thorny issue of nonlinearities that may confound invariance with respect to derived quantities like “tuning ratios” [Shera & Charaziak, 2019; see also Bergevin et al., 2015]. Further, there can be clear regions of elevated autocoherence in the “valleys” between magnitude peaks (e.g., 2.6-3 kHz in Fig.2C), potentially suggestive of coherent cooperative activity, just to a lesser extent than peak regions, amongst cellular generators.

Third, we took a comparative approach that showed that the timescale over which otocoherence can be maintained (as quantified by *N*_*ξ*_ in Fig.3) varies across species, with humans showing otocoherence time constants an order of magnitude longer than non-mammals. That the barn owl’s time constants were lower than humans can help explain why they can exhibit tall peaks that are surprisingly less likely to exhibit bimodal amplitude distributions [Bergevin et al., 2025]. Further, we showed *N*_*ξ*_/*π* appears to match well to the upper bound of ANF-derived *Q*_*ERB*_, indicating that SOAE activity can be used to infer the frequency selectivity of the auditory periphery. As illustrated by the spread of ANF-based *Q* values for a given characteristic frequency [CF, e.g., see Fig.4 in Shera et al., 2010 for chinchilla, or Fig.11 in Koppl, 1997 for barn owl], one might reasonably assume that similar-CF ANFs do not have a specific Q value. Rather, there is a range of *Q* values for the ANFs at any given CF when pooling data across animals/ears. Our data indicate that *N*_*ξ*_-based measures from SOAE activity are indicative of the upper bound of that range. That is, the mostly sharply tuned ANFs are the ones tied into regions of SOAE generation with strong self-coherence. The data shown in Fig.3B indicate that *N*_*ξ*_ values for Tokay gecko are slightly larger than those for the green anole (Fig.3B), consistent with the observations of longer auditory brainstem response (ABR) wave-I delays [Brittain-Powell et al., 2010], and suggestive of sharper tuning in gecko.

Lastly, we demonstrated different ways that the self-coherence can be calculated (i.e., time-delayed 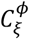 versus the frequency-neighbor referenced 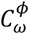) which might yield different insights into SOAE generation mechanisms (e.g., compare red versus blue/black curves in Figs.1C and 2C). While not systematically examined in this study, the 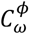 measure has different underlying biophysical motivations (e.g., generators can be spread across a narrow range of frequencies).

Applying these measures to study simulated waveforms from models as well as canonical examples (such as a noise-driven van der Pol oscillator) are natural next steps to ascertain any mutually exclusive information captured by either referencing strategy.

### Why is Otocoherence Lost?

To set the stage, we distinguish between intrinsic versus extrinsic noise, both in terms of physiological systems and (more generally) theory and simulation. Intrinsic noise would be that which influences a given system to affect a stochastic response from it. Extrinsic noise would be something additive to that response (e.g., microphone noise). Consider the NDDHO (Fig.3C) for example. There, the intrinsic noise would be the noisy forcing term that drives the oscillator. As a result, the system responds in a stochastic fashion, but with the system’s characteristics imprinted onto such. Generally speaking, averaging helps reduce the effects of extrinsic noise (e.g., magnitude averaging reveals structure in spectral magnitudes), whereas the autocoherence methods described here help ascertain the system’s underlying response to intrinsic noise.

We argue that the loss of otocoherence 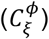 over time as shown in the colossograms of Fig.2 is primarily attributable to the role of intrinsic noise. Extrinsic, broadband additive noise would decrease the consistency of phase accumulations, but it should do so uniformly across timescales: it would add the same amount of randomness to each pair of phases, regardless of how far apart they are in time. The possibility of extrinsic non-additive (e.g., multiplicative) noise complicates this argument; however, the extreme differences across species largely preclude the possibility that the dominant factor is measurement noise. Therefore, we claim that the persistence and gradual loss of otocoherence is due—at least in part—to intrinsic noise affecting the phase stability of the underlying SOAE generation mechanism(s).

Our comparative work thereby indicates that some types of ears are less susceptible to intrinsic noise (e.g., Brownian motion due to fluid inner ear fluids), maintaining otocoherence over longer reference times *ξ* and thereby affecting tuning. Understanding precisely how and why will be important for auditory biophysics, but presumably the otocoherence revealed in “spontaneous” activity is closely related to the ear’s ability to actively respond to sound, especially at lower levels close to threshold. Given vast differences in morphology across ears exhibiting SOAE activity (e.g., anoles appear to lack basilar membrane traveling waves present in the mammalian cochleae), the methods and results described here should serve as a means to test the universality of mammalian-centric SOAE generation models such as the “‘active’ standing-wave model” [Shera, 2003].

### Looking Ahead & Next Steps

We believe the term “spontaneous” should be biophysically reserved to describe activity that is a result of the system’s functionality in the face of noise, rather than activity that is (otherwise uninteresting) noise itself. For example, ANF activity and its connection to ribbon synapse formation [Heil & Peterson, 2017; Coate et al., 2018]) could be considered spontaneous in this way: their stochastic nature leads to beneficial purpose. Broadly speaking, our study indicates that while SOAE appears to be noisy, it is actually indicative of how the system responds to sound. Pathological cases of SOAE will arise [e.g., Ruggero et al., 1983; Cheatham et al., 2016], but the fact remains that SOAE generation is a normal (albeit not universal) part of auditory function.

Typically, multiple waveforms were acquired from individual human and animal subjects, either over the same experimental session or different days. For humans, large amplitude peaks do appear relatively more stable (in peak frequency, magnitude, and otocoherence) across measurements, though quantification of this stability was beyond our current scope. However, some smaller subthreshold peaks (< 2 dB above surroundings) not included in Fig.4 analyses appear to vary more dynamically. Future work should examine whether these smaller peaks are spurious artifacts or represent transient/less coherent regions of SOAE generation.

A testable heuristic would be that human SOAE activity can be categorized as arrayed along a line. Each end would be designated as “stable”: one extreme representing moderate to large amplitude/coherent SOAE peaks, the other no peaks at all (i.e., quiescent). The middle would then represent a variable range of “unstable”, where smaller SOAE activity can dynamically change over various timescales, due to various factors such as thermal noise, efferent effects (i.e., top-down control), interactions between intracochlear generators, etc. Tonotopically, different frequency regions (or equivalently, spatial extents along the basilar membrane) would be characterized by placement along the line. One approach to elucidate such a framework could be a “coherogram”, where 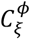 is calculated on “medium-sized” segments of the waveform (each containing enough smaller segments to calculate 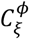) to explore the time-dependence of otocoherence. It could then be directly visualized in a fashion similar to a (magnitude-based) spectrogram and compared to transiently-evoked measures such SSOAEs. Generalizing beyond humans to other mammals, our new methods characterizing the timescale over which otocoherence is maintained can help distinguish subtle phenotypic effects in different mouse genotypes where the effects of mutations still remain poorly understood. For example, what does the increased incidence of SOAE activity in knockout mice where the TM is modified [Cheatham et al., 2016] tell us about disruptions to active high frequency (>10 kHz) cochlear mechanics?

Aside from further refinement of the methods described here (e.g., optimize choice of *ξ*-lag durations, introduce artifact rejection strategies) and expanding comparative analyses (e.g., systematic comparisons of otocoherence measures across disparate inner ear morphologies or variation in body temperature of ectotherms, such as in lizards), the approach could be adapted and applied to other types of empirical waveform data to study autocoherence measures of spontaneous activity. For example, auditory-evoked potentials [e.g., Franowicz & Barth, 1995], ANF firing patterns in the absence of external stimuli [Heil & Peterson, 2017], hair cell bundle oscillations affected by a weak applied force [e.g., Roongthumskul et al., 2013], and SOAE affected by transient stimuli that induce entrainment effects. Additionally, autocoherence measures could readily be computed from simulated waveforms from models of SOAE generation [e.g., Bowling et al., 2019] to benchmark them with respect to physiological SOAE activity. Even non-auditory measures such neural perisynaptic dynamics related to memory [Losi et al., 2025] could be explored under the proposed autocoherence framework.

Lastly, one ripe area for exploration is to characterize otocoherence between multiple spontaneous signals measured simultaneously where the generators may be linked in some fashion. As mentioned prior, the two ears of lizards are coupled by an interaural canal (IAC), allowing for weak acoustic coupling that apparently allows the inner ears to synchronize [e.g., Roongthumskul et al., 2019; Bergevin et al., 2020]. A “binaural otocoherence” defined similarly to 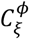 (where each pair of segments contains one drawn from each ear) could reveal causal relationships between the two ears. Moreover, it could take advantage of the fact that there would not be temporal overlap between segments, thereby reducing the need for (and potential limitations of) dynamic windowing [see Whiley et al., 2025].

## Materials and Methods

### Extracting Phase

We first introduce the notation 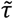 and 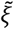 to denote the quantities *τ* and *ξ* in units of samples (i.e., 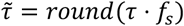), where *f*_*s*_ is the sampling frequency in Hz). The short-time Fourier Transform (STFT) provides estimates for the magnitude and phase of narrowband signal components as a function of both time and frequency [Oppenheim, 1999]. This is calculated by breaking the SOAE signal into a series of short windows and taking the DFT of each one. The STFT of our SOAE signal *x*[*n*] (with *n* indexing discrete time samples 1/*f*_*s*_ seconds apart) is

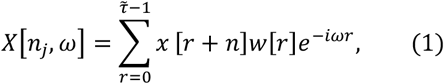

where we are using a window *w*[*n*] with 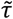 nonzero samples (*w*[*n*] is a “boxcar” window—i.e., *w*[*n*] = 1 for *n* = 0, 1, … 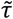 —unless otherwise indicated). We will evaluate at the standard DFT frequencies for a real signal *ω* = *ω*_*k*_ ≡ 2*πk*/*N, k* = 0,1, … *N*/2 (note *ω*_*k*_ is in radians per sample, corresponding to *f*_*k*_ = *ω*_*k*_ *f*_*s*_/2*π* in units of Hz) where the integer *N* (assumed to be even and ≥ 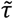) controls the amount of zero padding. The STFT is evaluated at evenly spaced points throughout the signal *n*_*j*_ ≡ *Hj, j* = 0,1, … (e.g., a hop size of *H* samples).

The STFT is a series of DFTs, each taking a ‘snapshot’ of the magnitude and phase of each frequency component at a given point in time. For example, the magnitudes squared *X*[*n*_*j*_, *ω*] ^2^ can be used to construct a spectrogram, tracking how the power spectrum changes over time. However, for our purposes it is illuminating to note that the STFT is also equivalent to a series of filtering operations. Specifically, 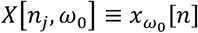 is simply the original signal after being filtered by a bandpass filter with impulse response 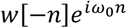 and frequency response *W*(*ω* − *ω*_0_), where *W*(*ω*) is the discrete-time Fourier Transform (DTFT) of the real window *w*[*n*] [Oppenheim, 1999]. Larger hop sizes correspond to sampling 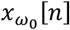 every *H* > 1 samples. We now have a set of narrowband signals 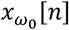 for a range of *ω*_0_. Note these are complex signals (since the filter *W*(*ω* − *ω*_0_) is asymmetric in frequency), from which we can take the complex argument to give a collection of phases as a function of both time and frequency *ϕ*[*n*_*j*_, *ω*] = arg(*X*[*n*_*j*_, *ω*]).

To extract information from these phases, our first method references each phase to the phase of another (nearby) frequency bin at the same point in time *Δϕ*_*ω*_[*n*_*j*_, *ω*] = *ϕ*[*n*_*j*_, *ω*] − *ϕ*[*n*_*j*_, *ω* + *Δω*]. Since each value *Δϕ*_*ω*_[*n*_*j*_, *ω*] represents information from both *ω*_*k*_ and *ω*_*k*+1_ in equal parts, all subsequent measures are plotted with respect to the average frequency (*ω*_*k*_ + *ω*_*k*+1_)/2. *Δϕ*_*ω*_[*n*_*j*_, *ω*] represents the difference in phase between one narrowband portion of the signal (bandpass filter centered around *ω*_0_) to another (centered around *ω*_0_ + *Δω*). For all figures, *Δω* was one DFT bin width 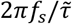. The second method references each phase to the phase within the same frequency bin, but at a point in time 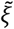 samples earlier. To achieve this, we calculate a 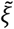 - shifted STFT *X*_*ξ*_[*n*_*j*_, *ω*] (defined identically as above except with a 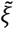 -shifted copy of the signal 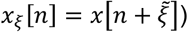 with resulting phases *ϕ*_*ξ*_[*n*_*j*_, *ω*] = arg(*X*_*ξ*_[*n*_*j*_, *ω*]). We can then define *Δϕ*_*ξ*_[*n*_*j*_, *ω*] ≡ *ϕ*_*ξ*_[*n*_*j*_, *ω*] − *ϕ*[*n*_*j*_, *ω*], an estimate for the accumulation in phase over 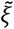 samples at frequency *ω*.

After subtracting the mean of each SOAE signal, STFTs (Eq.1) were calculated with *N* = 2^14^ to match the frequency grid used to pick the frequency centers (see The Characteristic Timescale *N*_*ξ*_). All analyses utilize a hop size *H* corresponding to 10 ms (*H* = 441 samples for *f*_*s*_ = 44100 Hz waveforms, *H* = 480 for *f*_*s*_ = 48000 Hz, and *H* = 500 for *f*_*s*_ = 50000 Hz). A hop size of *H* > 1 (for computational efficiency) is justified by the observation that when *H* << *τ* almost all samples in the window used to calculate a given phase estimate *ϕ*[*n*_*j*_, *ω*_0_] will also be used in the subsequent estimate *ϕ*[*n*_*j*+1_, *ω*_0_]. Since the same is true of the 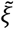 -shifted signal, the phase difference *Δϕ*_*ξ*_[*n*_*j*_, *ω*_0_] will be nearly identical to the subsequent *Δϕ*_*ξ*_[*n*_*j*+1_, *ω*_0_]; therefore, minimal information is lost by increasing *H* (up to a certain point, empirically determined for our methods to be ≈ 10 ms).

### Averaged Phase < |Δ*ϕ*| >

A relatively simple way to extract information from these phase differences is to wrap the *Δϕ* into the range [−*π, π*], take the absolute value, and then average across time points *n*_*j*_ for the quantity ⟨|*Δϕ*|⟩. For neighboring frequency referencing ⟨|*Δϕ*_*ω*_|⟩, this will track the tendency of neighboring narrowband components to be in phase versus antiphase with each other. For referencing in time, each phase difference *Δϕ*_*ξ*_ represents the accumulation in phase over *ξ* samples. Our goal is to quantify the consistency of this phase accumulation as a means of quantifying the self-coherence of the oscillations over a particular reference time *ξ*. We first note that a pure sinusoid will always evolve a fixed amount 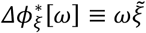 over this time. One option is to subtract off this “expected” accumulation, replacing each *Δϕ*_*ξ*_[*ω*] with 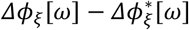. Then for frequencies where the oscillations are more consistently sinusoidal, each phase will be more “inphase” with the one 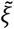 samples later 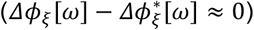, and the value of (|*Δϕ*|; _*ξ*_) will be lower (this is the approach used in Figure 1B).

### Auto-/otocoherence 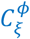

We can also directly track the self-coherence (or “autocoherence”) of the oscillations in a way which is agnostic to the particular value around which the phase differences may (or may not) tend to cluster. This is accomplished via a vector strength measure, which respects the periodicity of the phases in a more natural way; namely

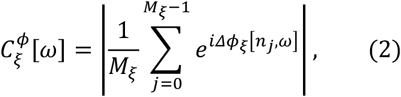

With 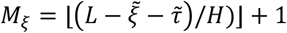 the maximum number of 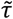 -length segments which can be extracted from *x*[*n*] and also have a full (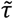 -length) reference segment 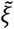 samples away (*L* = (length of *x*[*n*])). If the phase difference *Δϕ*_*ξ*_ remains relatively consistent, the resulting average vector will have a magnitude near 1, while if it fluctuates significantly the average vector will have a small magnitude closer to 0. This results in 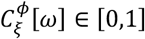, quantifying the self-coherence of the oscillations over the timescale *ξ* as a function of frequency; we refer to this as an autocoherence spectrum. An analogous measure for the frequency referenced phases 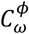 can also be defined by replacing *Δϕ* with *Δϕ* in the above definition for 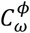. Again, the symbol 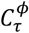 (and ⟨|*Δϕ_τ_* |⟩) refers to 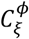 (and (*Δϕ_ξ_*)) in the special case that the reference time *ξ* equals the window length *τ*.

### Colossograms and the Intrinsic Gradient

We introduce the term “colossogram” to refer to a colormap displaying the full two-variable function 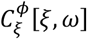, used to visualize how the decay in autocoherence over increasing reference time *ξ* depends on frequency *ω*. However, straightforward calculation of 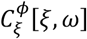 with a fixed window suffers from an “intrinsic gradient” of spurious autocoherence values. All frequencies (almost regardless of the nature of the signal) show high autocoherence at low *ξ* values, fading out as *ξ* is increased (see SI Fig.2A).

This intrinsic gradient is a consequence of the time-frequency tradeoff inherent to all filtering operations: bandpass filtering localizes in frequency and therefore smears timing information. This ubiquitous principle is seen directly in our case by considering the behavior of a stochastic process like white noise, which we would like our measure to quantify as perfectly “incoherent.”

Consider the calculation of 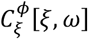 for a white noise signal over a short reference timescale relative to the window length (*ξ* << *τ*). The phase estimates arg(*X*[*n*_*j*_, *ω*]) will be scattered uniformly throughout [−*π, π*]. However, the segment used in the calculation of *X*[*n*_*j*_, *ω*] is nearly identical to the segment used in *X*_*ξ*_[*n*_*j*_, *ω*]: our stipulation *ξ* << *τ* means that they share nearly all samples, and so the sums in Eq.1 contain nearly identical terms (up to a phase factor 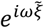 from each sample being shifted relative to the basis sinusoid). We then have the approximation 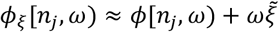, leading to very consistent lists of 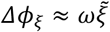 and therefore to spuriously high values of 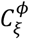 (and spuriously low values of < |Δ*ϕξ* | > since this approximation means 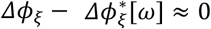 ; see Fig.1B). This argument explains why the spurious autocoherence is most pronounced for *ξ* << *τ* and no longer an issue for *ξ* ≥ *τ*. To address this, we implement a “dynamic windowing” method which has the visual effect of “peeling away” this gradient to reveal the structure underneath (see SI Fig.2C).

### Dynamic Windowing

We would like a single method that can be satisfactorily applied to the wide range of SOAE signals for cross-species comparison. To this end, a viable option to address the intrinsic gradient is to decrease *τ* (widening the bandwidth) only when necessary; namely, for short timescales *ξ*. The above discussion suggests simple approach: for all *ξ* less than some originally chosen *τ*_0_, use a modified window length *τ* = *ξ*. However, this means that as *ξ* increases, there is a point at which the modified *τ* abruptly stops increasing and becomes fixed at the originally chosen *τ*_0_. This transition manifests in the calculated autocoherence decay as an abrupt kink which interferes with our eventual goal of fitting a smooth exponential decay function.

Increasing *τ* without bound (i.e., *τ* = *ξ* always) would avoid this discontinuity, but (practical issues involving the finite lengths of our signals aside) this would mean approaching the scenario where the signal is filtered with an infinitely thin bandpass filter. In this limit, the filtered signal is a pure sinusoid with perfectly consistent phase accumulation and therefore trivially perfect autocoherence. Instead, we would like our dynamic windowing method to asymptotically approach some static window (with nonzero filtering bandwidth) in a smooth way. To achieve this, we keep *τ* fixed and instead multiply the original fixed window *w*_0_[*n*] by another window *w*_*ξ*_[*n*] which begins narrow and widens smoothly with *ξ*. A convenient choice for *w*_*ξ*_[*n*] is a Gaussian window with standard deviation (in units of samples) 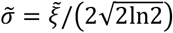 ; that is,

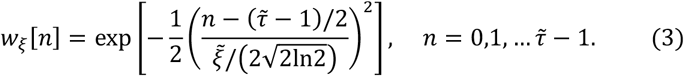

Since the full width at half maximum (FWHM) of a Gaussian window is 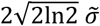, this choice of σ-results in the FWHM of the window being precisely equal to 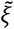. Then when 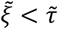 (the case where there are signal samples shared between a window and its *ξ*-advanced partner), the application of *w*_*ξ*_[*n*] applies weights to the shared samples that are always ≤ max(*w*_*ξ*_ [*n*])/2 in one of the two frames (see SI Fig.1). This “dynamic window” then reduces the intrinsic gradient significantly, while also retaining frequency resolution at higher *ξ* where the gradient is no longer prominent. If further reduction is desired, an additional parameter *ρ* can be introduced as 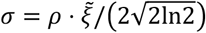 (so the Gaussian’s FWHM is 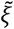), at the cost of widening the bandwidth even further for low *ξ*. For small *ξ*, the shape of the combined window *w*_*ξ*_[*n*] ⋅ *w*_0_[*n*] will be dominated by the narrow Gaussian *w*_*ξ*_[*n*], while as *ξ* → ∞ all coefficients of *w*_*ξ*_[*n*] approach unity and the combined window will asymptotically approach *w*_0_[*n*].

It remains to choose the asymptotic bandpass filter *w*_0_[*n*]. The shape of its frequency response depends on the window function *w*[*n*], with its length 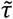 controlling the bandwidth. We chose the flat top window (as implemented in SciPy); this choice was motivated by the window’s relatively flat frequency response across the center (passing most of the SOAE activity region of interest in its original proportions) as well as its low sidelobes minimizing leakage from distant frequencies [D’Antona and Ferrero, 2006]. We quantify the width of this filter with the half-power bandwidth (twice the distance from *ω*_0_ at which the power gain of the filter falls to half). This bandwidth was chosen as 50 Hz, small enough to avoid ever including neighboring regions of SOAE activity. For the flattop window, this was achieved with 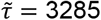 for *f*_*s*_ = 44100 Hz signals, 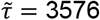 for *f*_*s*_ = 48000Hz, and 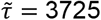 for *f*_*s*_ = 50000 Hz (each corresponding to *τ* ≈ 74.5 ms).

### The Characteristic Timescale *N*_*ξ*_

Once an SOAE frequency center is identified (see below), we would like to extract a characteristic timescale for its loss of autocoherence over increasing reference distance in time. Since nearly all decays were approximately exponential, we use the time constant *T*_*ξ*_ of an exponential function 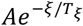. After calculating 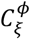 over a grid of *Δξ* = 1 ms, we fit this function (via nonlinear least squares) to the decay. For very small timescales *ξ*, our combined window *w*[*n*] = *w*_0_[*n*]*w*_*ξ*_[*n*] corresponds to a bandpass filter that is extremely wide. Initially, the narrowing of the bandwidth with increasing *ξ* caused by the dynamic windowing approach initially improves the autocoherence more so than it is degraded by the increasing timescale. We therefore start the fit shortly after it has begun to monotonically decline; specifically, we start the fit at the *ξ* for which the autocoherence has decreased to 90% of its peak value and continue until it reaches 10% of this peak value. This approach provided automatic identification of the relevant decay region.

For comparison with the nondimensionalized measure of tuning from auditory nerve fibers *Q*_erb_ [see Bergevin et al., 2015], we multiply each extracted time constant *T*_*ξ*_ by the center frequency *f*_0_ for a value *N*_*ξ*_ = *T*_*ξ*_ ⋅ *f*_0_. This then represents the timescale for loss of otocoherence in “number of cycles” rather than absolute time—that is, the number of cycles a sinusoid at frequency *f*_0_ would pass through over *T*_*ξ*_ seconds. These constants are plotted in Figure 3B after division by 2*π* to bring their scale into the range of *Q*_erb_.

For each species (human, anole, Tokay gecko, and barn owl), 60 second recordings from four subjects showing SOAE activity were used. For each subject, four SOAE frequency centers were chosen in a Welch-averaged power spectral density (PSD) spectrum [Welch, 1967], calculated with segments of length 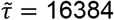, *H* = 8192 (50% overlap), and a Hann window. As much as possible, the selection of PSD peaks was chosen to have a wide range of frequencies, prominences, and proximities to other peaks. The frequency bins themselves representing the “frequency center” were then chosen simply as the maximum value in the peak region. In three cases, all from humans, the decays at a particular frequency center appeared poorly fit by an exponential function. Each was replaced by another picked in the same way as the original, and in all cases the first replacement had a satisfactorily exponential decay.

### Incorporating Magnitudes for 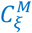

Finally, we define a variation of autocoherence which incorporates the magnitudes.

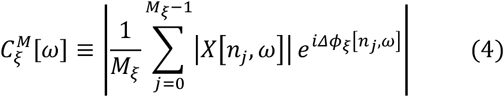

Peaks in this new spectrum then represent frequencies that were relatively high in both magnitude and phase autocoherence. The analysis displayed in Fig.4B-E was performed on twelve human SOAE waveforms recorded at 44100 Hz. Before the spectra were calculated (and peaks identified via the 2 dB criterion described in Results), waveforms were cropped to regions that appeared relatively artifact-free to prevent identification of peaks that were unrelated to SOAE activity. For 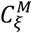, the parameters 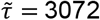 samples (∼69.66 ms) and 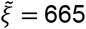 (∼15.08 ms) were used, along with the window *w*_*ξ*_[*n*] as defined in Eq.4 was implemented (with no “asymptotic” window, i.e., *w*_0_[*n*] = 1 for *n* = 0, 1, … 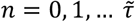).

Averaged magnitude spectra were calculated as a simple average over the magnitude of all segments 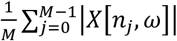, where *M* was the number of segments of length 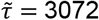 samples (∼69.66 ms, same as 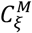) that could be extracted from the waveform (with *H* = 441 samples) and *w*[*n*] in Eq.1 was a Hann window. The choice to average unsquared magnitudes *X*[*n*_*j*_, *ω*] (rather than the modulus squared *X*[*n*, *ω*]^2^ typically used in Welch spectral averaging) is motivated by its relative resistance to outlier segments. Specifically, for a list of nonnegative real numbers the result of averaging their squares is always greater than (or equal to) the result of averaging the original list (and then taking the square), and the difference is precisely the variance of the list.

For SOAE signals, these outlier segments often represent artifacts during the measurement process which we would not want to identify as SOAE peaks. Finally, note that no scaling constants were utilized, so all figures depict both averaged magnitudes and 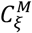 on a “floating” dB scale (i.e., the vertical offset is not meaningful).

### Noise-Driven Damped Harmonic Oscillator Simulations

A damped harmonic oscillator driven by standard normal white Gaussian noise was simulated via a method taking advantage of the linearity of the equation and the Gaussianity of the drive term to perform an “exact” integration [Nørrelykke & Flyvbjerg, 2011]. Ten 60s signals (each with a unique random drive term) were generated at *f*_*s*_ = 44100 Hz for each combination of *Q* ∈ [25, 30, 35, 40, 45, 50] and damped natural frequency *f*_*d*_ ∈ [1000, 2000, 3000] Hz (chosen to lead to PSD peak widths roughly on order with those observed in the SOAE signals). Nondimensionalized decay constants *N*_*ξ*_ were extracted using the dynamic windowing method with the same parameters as the (*f*_*s*_ = 44100 Hz) SOAE signals.

### Code

Analysis code can be found here, utilizing our Python package phaseco for autocoherence calculations.

## Acknowledgments

We gratefully acknowledge Yuttana Roongthumskul and Christine Koppel for sharing Tokay gecko SOAE data and owl/gecko auditory nerve fiber data respectively. CB, RW, and SP were supported by Natural Sciences and Engineering Research Council of Canada (NSERC) Grant RGPIN-430761-2013. NM was supported by NSERC Grant RGPIN-687216 (with early career supplement 675248), and an NSERC Canada research chair (693206). VV was supported by Czech Science Foundation grant 23-07621J. SP was also supported by the Fields Institute Undergraduate Summer Research Program (FUSRP). Comments from Julien Meaud, Yuttana Roongthumskul, and Christopher Shera are also gratefully acknowledged.

