## Supplemental Information for "Spontaneous otocoherence of the active ear"

### Spontaneous otocoherence of the active ear
### Supplemental Information

Seth N.S. Peacock, Václav Vencovský, Rebecca E. Whiley,
Natasha Mhatre, Christopher Bergevin

#### 1 SI: Comparison with Other Methods

##### 1.1 Narrowband Signal Decomposition

Note there are other methods of extracting narrowband representations of the signal for phase estimates as a function of time and frequency, such as the wavelet transform or by constructing the analytic signal after bandpass filtering. However, while these are useful alternative frameworks, they are ultimately equivalent to the STFT approach with the appropriate choices of parameters (Bruns 2004). For example, the wavelet transform uses a frequency resolution (or bandwidth of the equivalent bandpass filter) which is consistent across frequency in number of cycles rather than in absolute Hz. This can be applied in the STFT framework by simply using different values of $\tau$ for different frequencies (this approach is taken in Fransen, Ede, and Maris 2015). However, this would not be appropriate for our purposes in SOAE analysis, as the PSD bandwidths of SOAE peaks are relatively constant across frequency. When selecting out a portion of SOAE activity to quantify its phase autocoherence, it is therefore not clear why the width of such a portion should vary with frequency.

##### 1.2 Power Weighted $C_\xi^P$

Our definition of $C_\xi^\phi[\omega]$ treats each $\Delta\phi$ equally in an unweighted average. However, we discuss the following "power weighted" variant of the measure, equivalent (up to the method of obtaining the magnitude and phase as a function of time and frequency and the choice of whether or not to square the result) to the one used in Zhou and Dagle 2014 and Fransen, Ede, and Maris 2015. Even if phase is the primary object of interest, it may still be desirable to incorporate the magnitudes of each segment corresponding to $\Delta\phi$ as weights: if the magnitude of the signal component of interest fluctuates relative to a fixed noise floor, phase differences that are weighted higher will be those from segments with a higher signal-to-noise ratio (SNR) at that frequency and are therefore ostensibly more accurate phase estimates (Bruña, Maestú, and Pereda 2018). Each $\Delta\phi_\xi$ comes from two segments, and so a natural choice is to weight each one by the product of the two segments' magnitudes. This leads to an average over $|X_\xi[n_j,\omega)||X[n_j,\omega)|e^{i\Delta\phi_\xi[n_j,\omega)} = X_\xi[n_j,\omega]\overline{X[n_j,\omega]}$ (where $\overline{X}$ denotes complex conjugation). The result is then normalized to $[0,1]$, by the geometric mean of Welch estimates of the power spectral densities of $x[n]$ and $x_\xi[n]$ for the "power weighted" autocoherence

$$C_\xi^P[\omega] \equiv \frac{\left|\frac{1}{M_\xi}\sum_{j=0}^{M_\xi-1}\overline{X[n_j,\omega)}X_\xi[n_j,\omega)\right|}{\sqrt{\left(\frac{1}{M_\xi}\sum_{j=0}^{M_\xi-1}|X[n_j,\omega)|^2\right)\left(\frac{1}{M_\xi}\sum_{j=0}^{M_\xi-1}|X_\xi[n_j,\omega)|^2\right)}}$$

We are hesitant to concur with a claim regarding the amplitude dependence of this version of the measure made in Fransen, Ede, and Maris 2015. They acknowledge that this method is not truly "amplitude-independent" in the presence of noise, due to the effect of changing SNR across segments due to amplitude changes in the noise and/or signal across time. However, they claim that this amplitude independence holds "in cases where the signal is noise-free." Consider such a noise-free signal with oscillations vacillating between periods of high amplitude but low phase autocoherence (low "rhythmicity") and low amplitude but high phase autocoherence. Since the terms in the sum are weighted by the amplitudes of the two segments, it is clear that the high amplitude segments would dominate and the overall autocoherence of the signal would be biased towards that of the higher amplitude segments. We therefore use the more amplitude-independent measure of $C_\xi^\phi$, which is equivalent to $C_\xi^P$ with the amplitude of each segment normalized to 1. (For SOAE signals, the empirical difference is very small.) Nevertheless, we acknowledge that amplitude modulation may still affect even our amplitude normalized method by interfering with underlying estimates of the phases $\phi[n]$.

##### 1.3 Relationship of Two-Signal Coherence to $C_\xi^\phi$ and $C_\xi^P$

The power-weighted $C_\xi^P$ is then precisely equivalent to the standard Welch estimate of coherence between the two signals $x[n]$ and $x_\xi[n]$ (e.g. as implemented by SciPy, see Stoica, Moses, et al. 2005, Welch 2003), a relationship manifest in the presentation of past work (Zhou and Dagle 2014, Fransen, Ede, and Maris 2015). However, while this can be a useful analogy, we point out that the theory behind the measure of coherence breaks down in the particular case where one signal is a shifted copy of the first. The underlying theoretical coherence value the Welch method estimates is

$$\gamma(\omega) = \frac{|\mathbb{E}\left[X(\omega)Y^*(\omega)\right]|}{\sqrt{\mathbb{E}[|X(\omega)|^2]\mathbb{E}[|Y(\omega)|^2}}, \tag{1}$$

where $\mathbb{E}$ indicates a population expectation over all possible realizations of random processes $x[n]$, $y[n]$ and $X$, $Y$ are their infinite length DTFTs. When $y[n] = x[n+\xi]$, the DTFTs for any given pair of realizations at any given frequency $\omega$ are related via $Y = e^{i\xi\omega}X$, and so we have the trivial result $\gamma(\omega) = 1 \;\; \forall\omega$.

##### 1.4 Relationship of Autocorrelation to $C_\xi^\phi$ and $C_\xi^P$

A more apt comparison is to autocorrelation; specifically, $C_\xi^P[\omega]$ is approximately the magnitude of the unbiased sample autocorrelation of the narrowband filtered signal $x_{\omega_0}[n] \equiv X(\omega_0, n)$ at lag $\xi$. Assuming our signals are zero mean or have already had their means subtracted, the sample autocorrelation of $x_{\omega_0}[n]$ for lag $\xi \geq 0$ is (Brockwell and Davis 2002)

$$\hat{r}_{\omega_0}(\xi) = \frac{\frac{1}{M_\xi}\sum_{n=0}^{M_\xi-1} x_{\omega_0}[n]\overline{x_{\omega_0}[n+\xi]}}{\frac{1}{L}\sum_{n=0}^{n=L-1}|x_{\omega_0}[n]|^2}$$

where $L$ = length of $x_{\omega_0}[n]$. Under the mild assumption that

$$\sqrt{\left(\frac{1}{M_\xi}\sum_{n=0}^{M_\xi-1}|x_{\omega_0}[n]|^2\right)\left(\frac{1}{M_\xi}\sum_{n=0}^{M_\xi-1}|x_{\omega_0}[n+\xi]|^2\right)} \approx \frac{1}{L}\sum_{n=0}^{L-1}|x_{\omega_0}[n]|^2, \tag{2}$$

the following approximation holds.

$$C_\xi^P[\omega] \approx |\hat{r}_{\omega_0}(\xi)| \tag{3}$$

Similarly, if no dynamic windowing is employed, the non-power-weighted $C_\xi^\phi$ used in our main results can be interpreted as the magnitude of the sample autocorrelation of the narrowband signal after being amplitude normalized sample-by-sample, $x_{\omega_0}[n]/|x_{\omega_0}[n]|$ (note the denominator in the definition of $\hat{r}_{\omega_0}$ is then equal to 1).

However, this analogy ultimately breaks down as well. The first reason is the dynamic windowing method in which the bandpass filter applied to the signal is narrowed with each increasing autocorrelation lag $\xi$. Second, this definition of the autocorrelation as a single variable function assumes that the signal is wide sense stationary (WSS), a property which SOAE signals do not have (Haggerty, Lusted, and Morton 1993). Despite that, the analogy between autocorrelation and $C_\xi^\phi$ is a useful when paired with the fact that (for WSS signals where both are well defined and single variable functions) the PSD is the Fourier transform of the autocorrelation. Since most SOAE PSD peaks are roughly Lorentzian—a peak which is the Fourier pair of an exponential decay—it should not be surprising that the autocoherence decays are roughly exponential. Nor should it be surprising that—for a static windowing approach—the decay constants are inversely proportional to the width of a Lorentzian peak fitted to the filtered PSD (see Figure 3).

This provides a useful viewpoint from which to understand the "intrinsic gradient" imposed by the window function. Even if your input is flat white noise, the resulting spectrum after filtering with a bandpass filter will be approximately peak-like; the thinner the bandwidth, the more it will resemble a narrow peak, leading to longer and longer spurious decays. The window function imposes a structure on the filtered signal, a structure which can obscure the true underlying properties of the signal we wish to

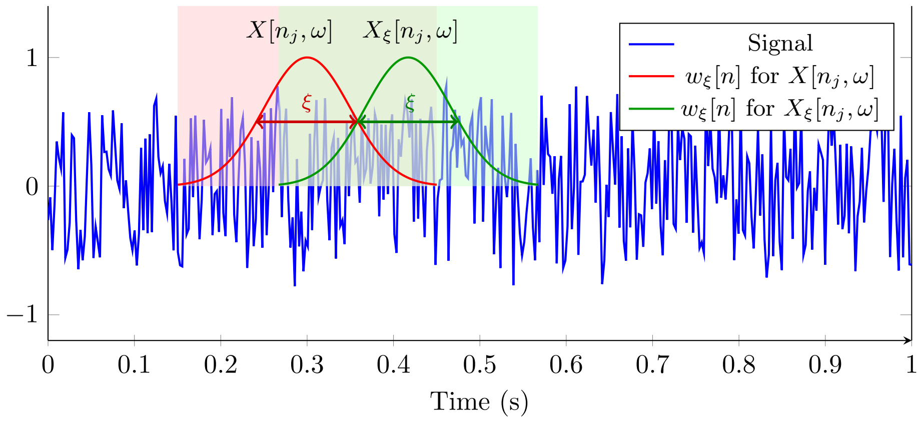


Figure 1: Diagram of the dynamic windowing approach for segment $j$. The dynamically adjusted Gaussian window $w_\xi[n]$ used in the initial segment is shown in red, while the Gaussian window used for its $\xi$-advanced partner is shown in green. Note the half width half maximum of the Gaussian windows are exactly equivalent to $\xi$ ($\rho = 1$), meaning that any samples shared between both frames receive weights from $w_\xi[n]$ that are $\leq \frac{\max(w_\xi[n])}{2} = \frac{1}{2}$. Since the asymptotic window $w_0[n]$ (not pictured) is also $\leq 1$, the ultimate window $w[n] = w_0[n] \cdot w_\xi[n]$ weights these shared samples by $\leq \frac{\max w[n]}{2} = \frac{1}{2}$.

extract. This further motivates the use of dynamic windowing; rather than choosing a single structure to impose on the signal, we sweep through a range of structures (the increasing width of the Gaussian window) and extract a single constant $N_\xi$ representing the signal's response to this range. This approach, alongside the nonstationarity of SOAE signals, provides the opportunity to extract information beyond that of a simple peak width in magnitude based spectra like the PSD (or rather, the spectrum obtained through a standard Welch estimate of PSD when stationarity is assumed to hold approximately).

#### 2 SI: Static vs Dynamic Windowing

Figure 4 displays the difficulty in choosing a single bandwidth (and therefore $\tau$) for SOAE analysis across all species. As discussed in **??**, we decided that varying $\tau$ across species would ultimately be undesirable for cross-species analysis, leading to the development of the dynamic windowing method. However, to have a frame of reference for this method, we performed a rough analysis with a static windowing approach. For each species, a $\tau$ was chosen based on an approximate assessment of an appropriate bandwidth based on the species' typical PSD peak width. This was chosen as 50 Hz for humans, 150 Hz for tokay geckos and anoles, and 300 Hz for owls. Outside of the use of a static flattop window, the decay constants were calculated using the same parameters as the main dynamic windowing analysis, with the exception of using a grid of $\xi$ spaced $\Delta\xi = 0.5$ms apart rather than 1ms to capture the quicker decays manifest with this approach.

Figure 3 compares the $N_\xi$ extracted from the two approaches. A significant amount of frequency centers (six from anoles, four from owls, three from tokays, and two from humans) had rather non-exponential decays with this static approach; these were excluded from the plot for a fair comparison of the methods. Note the structure in the dynamic windowing plot that is much less apparent in the static windowing plot. For the static windowing, we can compare the phase-based decay constants at $\omega_0$ to the width of the peak in the magnitude-based PSD spectrum of the filtered signal $x_{\omega_0}[n]$. (No such comparison is available for the dynamic windowing, since the filtered PSD would change with $\xi$.) As discussed in the previous section, this should not be surprising due to the relationship between the autocorrelation and the PSD of the signal $x_{\omega_0}[n]$ (at least, under the—at most approximately true for our signals—assumption of wide sense stationarity). In total, this figure suggests that the static windowing approach reveals information that is roughly proportional to information contained in the PSD (or rather, the PSD after the static window has been imposed on it), while the dynamically windowed autocoherence extracts different information.

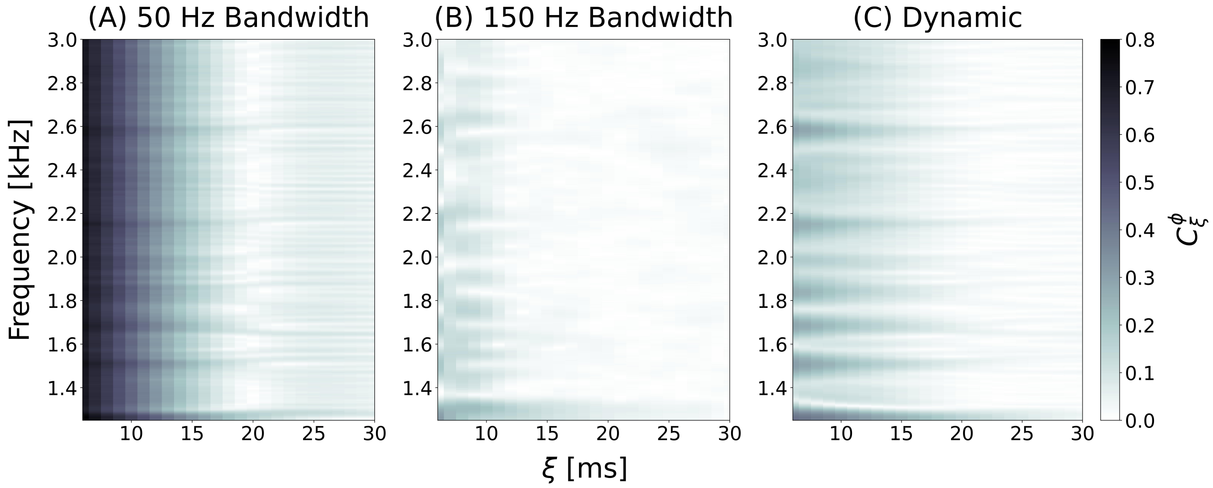


Figure 2: Comparison of windowing methods for an anole SOAE; A uses a 50 Hz static flattop window, B a 150 Hz static flattop window, and C dynamic windowing (with a 50 Hz flattop asymptotic window). The narrow static window has an obscuring intrinsic gradient, which the dynamic windowing method removes and reveals features underneath. The broad static window also removes the gradient, but heavily distorts the underlying features.

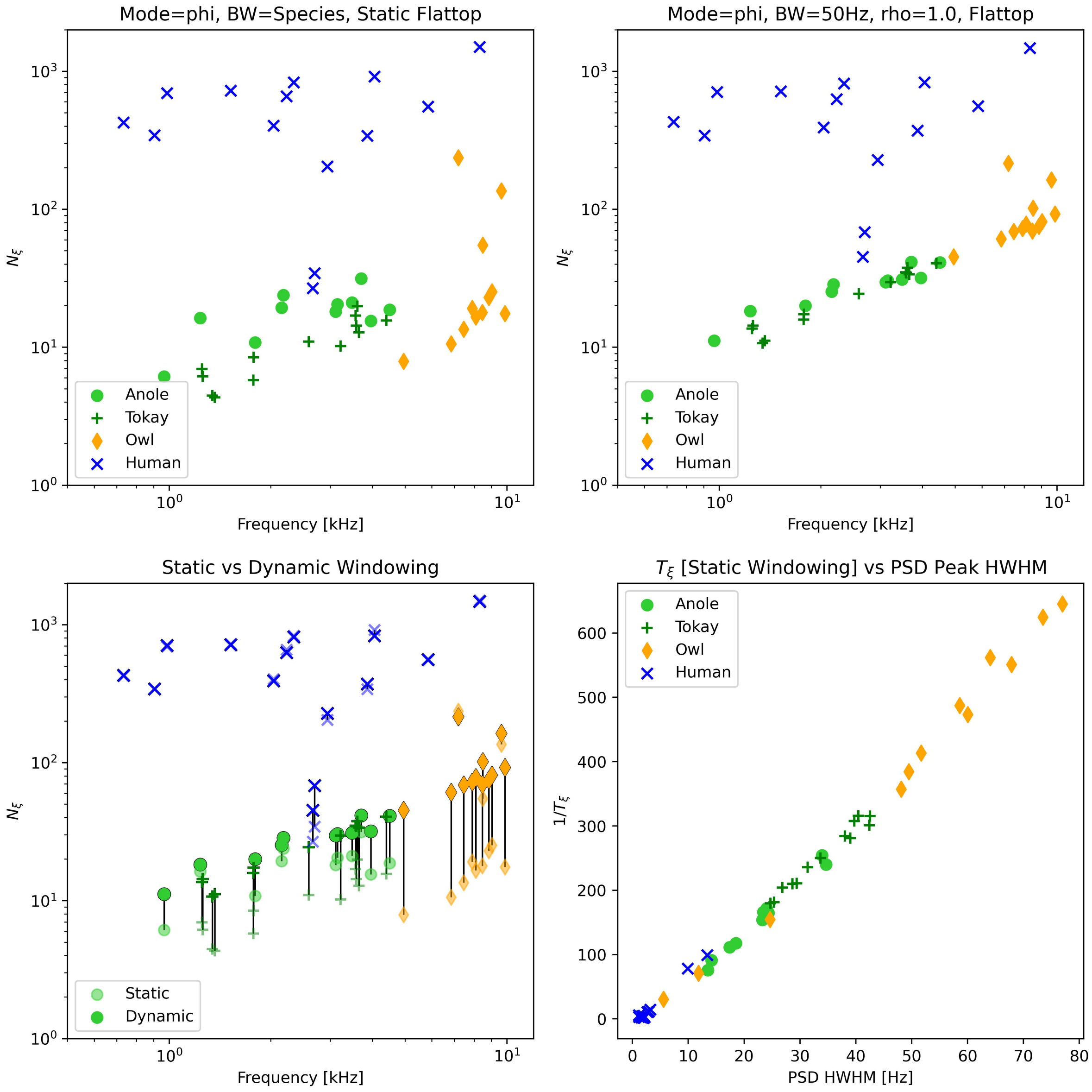


Figure 3: Comparison of the nondimensionalized decay constants $N_\xi$ extracted with static windowing (upper left) vs dynamic windowing (upper right). Frequency centers with significantly non-exponential decays when using static windowing were excluded from all four plots for a fair comparison of the methods. The lower left shows both on the same plot, with the static windowed constants faded. The lower right shows the direct proportionality of the inverse of the decay constants extracted via static windowing $(1/t(\xi))$ against the estimated half-width-half-maximum (HWHM) of the filtered PSD peak. For the latter, a Welch estimate of the PSD of the filtered signal $x_{\omega_0}[n]$ was calculated with a Hann window of length $\tau = 2^{14}$ and a hop of $\tau/2$, to which a Lorentzian peak was fitted and the HWHM extracted as the Lorentzian parameter $\gamma$.

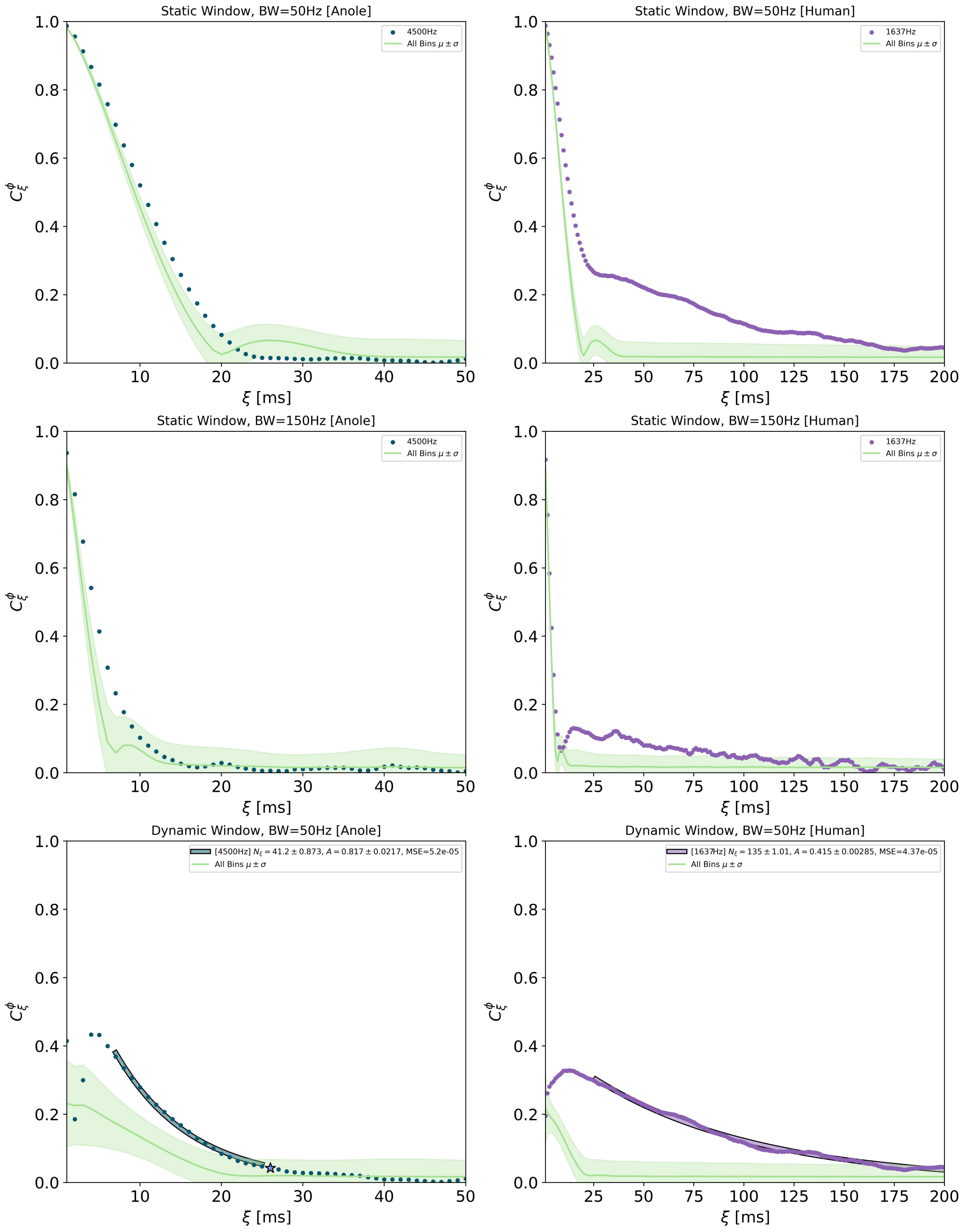


Figure 4: A demonstration of the difficulty in choosing a single static window for SOAE signals from all species. For this pair of SOAE frequency centers (anole in left column and human in right), we start with a static window with a bandwidth of 50Hz. At this bandwidth, the anole's decay is all but indistinguishable from the intrinsic gradient (approximated here by the mean ± standard deviation over all frequency bins). If we increase the bandwidth to 150Hz, this pushes back the intrinsic gradient enough that it may be reasonable to fit and interpret the anole decay. However, the human decay is squashed down and oscillates wildly at this wider bandwidth. This can be understood by noting that a 150 Hz bandwidth extends significantly beyond the width of a typical human SOAE PSD peak, so such a bandwidth band of coherent activity we wish to examine. With dynamic windowing, both frequency centers remain significantly above the intrinsic gradient throughout the decay and can be well fit by an exponential (the human still oscillates, but no more than the original static 50Hz window.)
